# All-Atom Structure Tokenizer: AtomLM-9b for Biomolecular Structure Prediction

**DOI:** 10.64898/2026.11.03.763913

**Authors:** Jiaxian Yan, Dongchu Xie, Zhicai Wang, Mingyang Xu, Zaixi Zhang

**Author notes:** Equal contribution (Jiaxian Yan, Dongchu Xie).

## Abstract

We develop an All-Atom Structure Tokenizer that compresses biomolecular geometry into discrete structural tokens and achieves high-accuracy all-atom reconstruction. A hierarchical geometric encoder and finite scalar quantization map chemical units to a shared vocabulary of 8,192 tokens, while a joint coordinate de-tokenizer reconstructs backbone, side-chain, nucleic-acid, and ligand heavy atoms. The tokenizer achieves all-atom lDDT scores of 0.832 and 0.835 on CASP14 and CASP15, respectively. Building on this tokenizer, we train AtomLM-9b, an autoregressive language model based on Qwen3.5-9B that predicts structural tokens from molecular identities and connectivity and decodes them into complete atomic structures. We demonstrate, for the first time, protein structure prediction performance comparable to AlphaFold 3 with an all-atom-tokenizer-based structural language model: on the FoldBench protein–protein task, AtomLM-9b achieves a success rate of 75.01%, compared with the published AlphaFold 3 result of 72.93%. These results establish discrete all-atom tokenization as an effective interface between high-fidelity atomic reconstruction and language-model-based structure prediction.

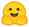huggingface.co/Scinetics-ai/all-atom-structure-tokenizer

## 1 Introduction

Accurate biomolecular modeling requires both global organization and local chemical precision. Protein folds, side-chain packing, ligand poses, and nucleic-acid interfaces are coupled, yet literal atom-by-atom representations can make autoregressive generation expensive. A structural vocabulary offers an alternative: compress geometry into discrete symbols, model their dependencies with a large sequence model, and recover coordinates with a generative de-tokenizer.

The All-Atom Structure Tokenizer provides this interface at chemical-unit granularity. Its encoder contextualizes the complete assembly before quantization; a shared vocabulary represents residues, nucleotides, and ligand atoms; and a joint de-tokenizer recovers atomic geometry with inter-unit refinement. AtomLM-9b is an autoregressive language model that learns the resulting structural-token sequences using a chemistry-conditioned Qwen3.5-9B backbone [3]. This division allows discrete sequence modeling and continuous geometry reconstruction to be optimized and evaluated separately. We achieve high-accuracy all-atom reconstruction and establish protein structure prediction performance comparable to AlphaFold 3 with the resulting structural language model.

### A shared vocabulary, different chemical units

The structural vocabulary is shared across molecular types, but the unit definition follows chemistry. A protein residue includes backbone and side-chain atoms; a nucleotide includes base, sugar, and phosphate; a small-molecule unit contains one heavy atom. Figure 1a shows ubiquitin from PDB 1UBQ and the XK2 ligand from 1HVR as examples of observed geometries.

**Figure 1.**
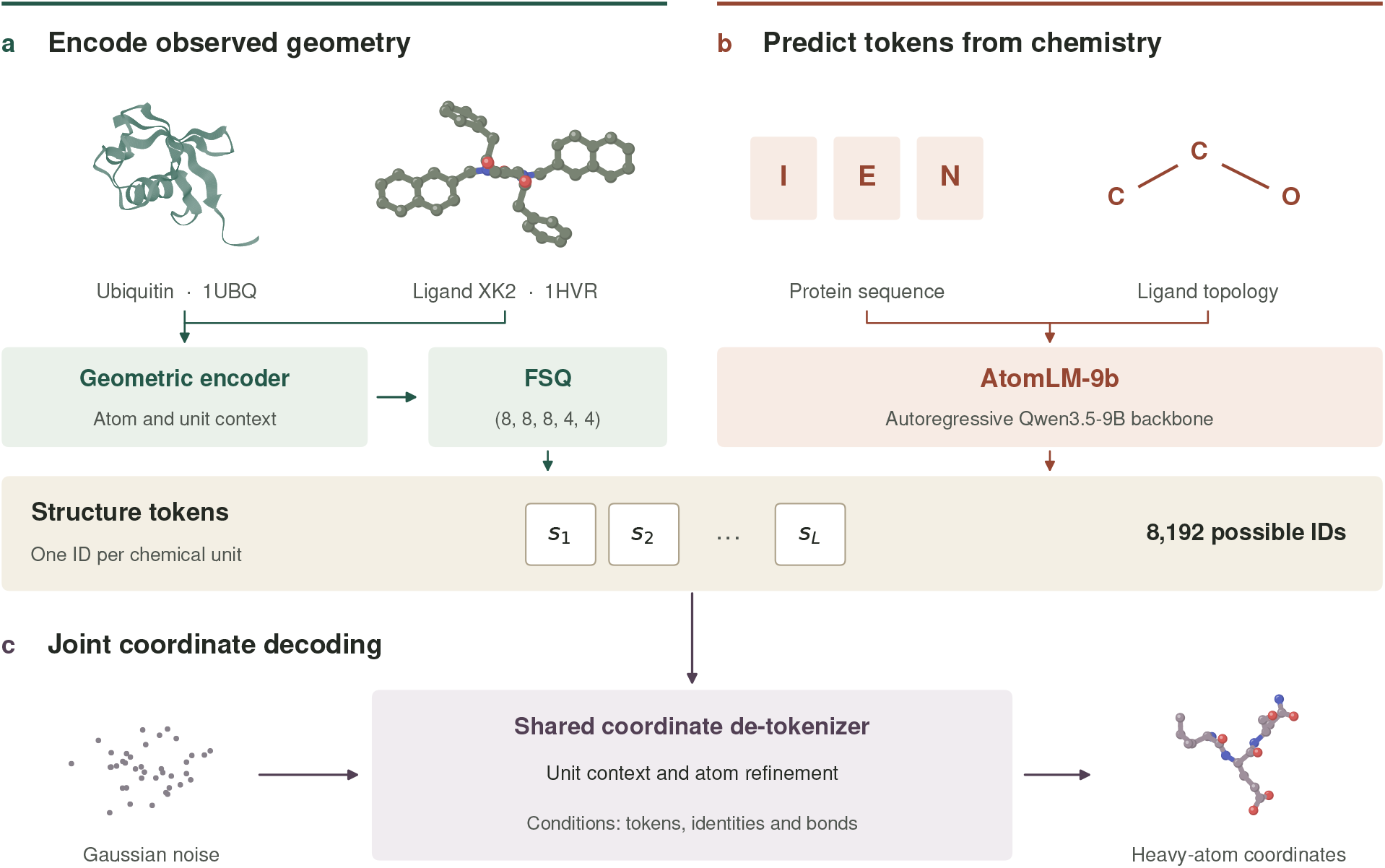
Two routes to all-atom structure tokens. (a) The tokenizer encodes observed geometry and quantizes one feature per chemical unit. (b) AtomLM-9b predicts tokens in the same vocabulary from molecular identities and bonds. (c) Both routes use the same de-tokenizer to recover heavy-atom coordinates from tokens and chemistry. Molecular views illustrate experimental structures.

### A discrete bottleneck with atomic reconstruction

The discrete ID is the only geometric information passed from encoder to de-tokenizer. Chemical identities and connectivity remain separate conditioning inputs. Five-dimensional FSQ with levels (8, 8, 8, 4, 4) provides a vocabulary of 8,192 IDs. The de-tokenizer expands unit context into atom features and refines the full assembly jointly, retaining atomic output resolution without assigning a separate token to every protein atom. At inference, the language model supplies tokens from chemistry alone.

### Relation to prior work

Hierarchical atom–unit encoding draws on interaction representation learning in ATOMICA [1]. FSQ supplies a fixed discrete space without learned nearest-neighbor codebook updates [2]. The All-Atom Diffusion Tokenizer (ADT) uses diffusion to reconstruct protein backbone and side-chain coordinates from one structural token per residue [8]. Our system design combines assembly-level contextualization, multi-level quantization, inter-unit atom refinement, and a 9B chemistry-conditioned language model.

#### Scope

*All-atom* denotes explicit modeling of all supported heavy atoms, including protein side chains. The chemical interface covers proteins, nucleic acids, small molecules, and their complexes. Hydrogens, solvent, and unmodeled atoms are outside the primary metrics.

## 2 A unified all-atom tokenizer

### 2.1 Chemical units, atom identities, and topology

Let a molecular assembly be *M* = (*C, B, X, m*), where *C* contains molecular and atom identities, *B* is the covalent bond graph, *X* ∈ ℝ^*N×*3^ contains heavy-atom coordinates, and *m* marks observed coordinates. A partition 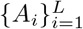 assigns atoms to structural units. Chemistry determines unit membership and the expected atom inventory. Observed coordinates define the encoder graph during tokenizer training, and the mask gates reconstruction losses. Native observation masks are not supplied to the language model or de-tokenizer at prediction time.

Protein residues provide compact units with explicit side-chain membership. Small molecules use atom units to avoid ambiguity from variable motif decompositions. The shared vocabulary therefore compresses protein sequences while preserving an unambiguous ligand atom ordering. Standard residue templates and Chemical Component Dictionary (CCD) atom names preserve correspondence; unsupported components are rejected or represented by an explicit atom-level fallback.

### 2.2 Hierarchical geometric encoding

The encoder separates local chemical geometry from assembly-level organization. Eight equivariant atom blocks first exchange messages over typed geometric edges. Learned pooling then maps atom states to chemical units, and four unit-context layers propagate information across the complete assembly. A rotation-invariant readout produces one 512-dimensional feature per unit for FSQ. Figure 2 details this hierarchy.

**Figure 2.**
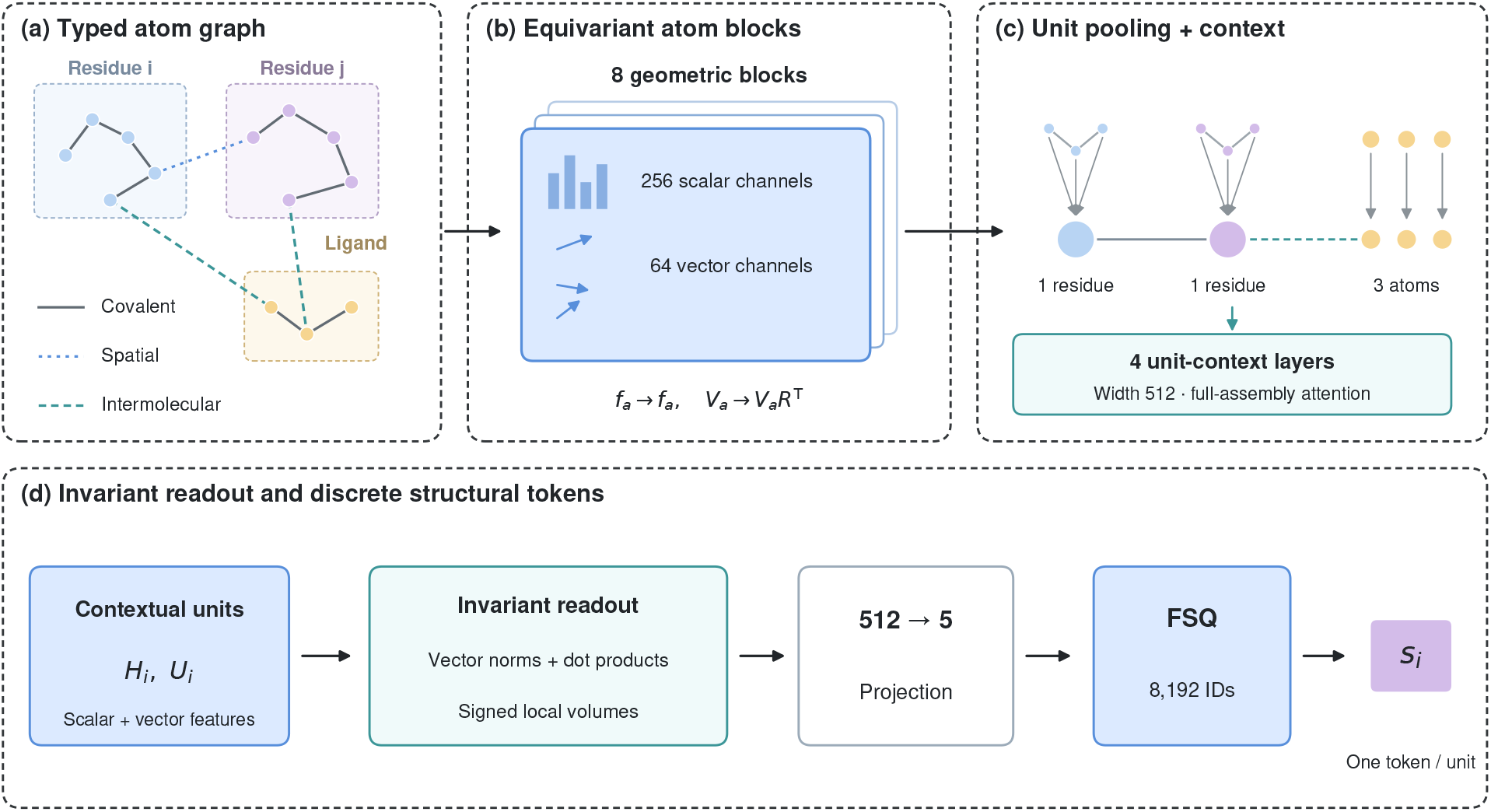
Hierarchical geometric encoder. (a) Covalent, intramolecular spatial, and intermolecular edges carry distinct message types. (b) Atom blocks update scalar and directional features. (c) Residue atoms pool into one unit; ligand atoms retain one-to-one units. Unit attention exchanges assembly context. (d) Invariant contractions and signed local volumes feed a 512-to-5 projection and FSQ. Graph layouts and node counts are schematic.

#### Typed neighborhoods and atom states

Covalent bonds are always retained, including cross-chain bonds. Additional neighbors comprise up to 32 intramolecular and 16 intermolecular atoms within 12 Å; these spatial caps do not remove covalent edges. Edge features *e*_*ab*_ combine bond type, relation type, and radial distance encodings. Atom states contain 256 scalar channels *f*_*a*_ and 64 vector channels *V*_*a*_ ∈ ℝ^64*×*3^. Scalar inputs encode element, charge, aromaticity, atom role, chemical-unit identity, and chain membership. Relative directions *r*_*ab*_ supply orientation without using an absolute coordinate frame.

#### Equivariant atom updates

A scalar–vector message block uses invariant contractions to parameterize scalar messages and vector gates [1]:

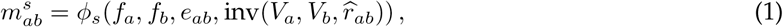

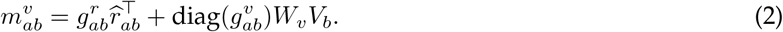

The gates 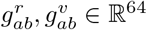 are learned from invariant edge features. Neighbor messages are summed and applied through residual updates. Scalar normalization and vector-norm gating preserve rotation covariance; the eight blocks share the feature dimensions, with separate layer parameters.

#### Atom-to-unit pooling

Invariant attention weights aggregate atoms within the chemistry-defined partition:

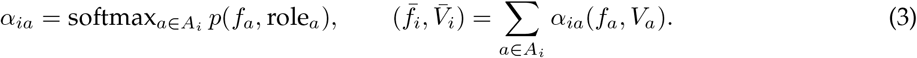

Protein residues and nucleotides pool their modeled heavy atoms. A ligand atom forms a singleton unit, so its pooling weight is one. Empty observed units are excluded from tokenizer reconstruction. The expected prediction inventory remains chemistry-defined. Pooling weights are invariant; vector averages therefore remain equivariant.

#### Assembly-level unit context

Scalar unit features are projected to width 512. Four attention layers use chain-relative positions, unit connectivity, and invariant geometric biases from unit centroids and vector– direction contractions. Contact types are inherited from atom edges. Global attention communicates beyond the atom cutoff, allowing a residue token to reflect both internal conformation and interface context. Vector unit features are aggregated with invariant attention weights, preserving their transformation rule. The resulting scalar and vector unit states are denoted *H*_*i*_ and *U*_*i*_.

#### Invariant, chirality-sensitive readout

The final feature *h*_*i*_ ∈ ℝ^512^ combines scalar context, vector norms, selected inner products, and signed local volumes. For chemically ordered neighbor triplets, 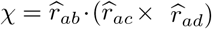 preserves handedness under proper rotations; degenerate triplets are masked. These features remove translation and orientation dependence while retaining reflection sensitivity. Only quantized IDs, rather than continuous encoder states, reach the coordinate de-tokenizer.

### 2.3 Finite scalar quantization

We project each unit to *d*_*q*_ = 5 bounded scalars and use levels *l* = (8, 8, 8, 4, 4). The parameterization and mixed-radix ID mapping are

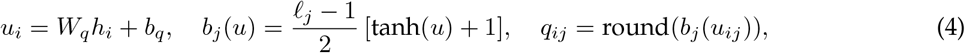

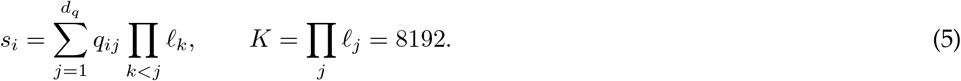

Straight-through gradients approximate differentiation through rounding. Quantized coordinates are rescaled to [−1, 1] before de-tokenizer projection. The allocation of levels across five dimensions defines the discrete geometry interface; a capacity-matched binary variant is evaluated in the ablation study. An 8,192-way token carries at most 13 bits, regardless of hidden width. FSQ avoids learned-codebook updates.

## 3 Conditional all-atom de-tokenization

### 3.1 From chemical units to atoms

The de-tokenizer models *p*_*ϕ*_(*X* | *S, C, B*), with *S* = (*s*_1_, …, *s*_*L*_). It receives discrete IDs, the full chemical graph, and noise. No continuous encoder states, native distances, native local frames, or coordinates bypass the quantizer. This makes reconstruction a genuine test of the discrete interface.

A unit-level diffusion transformer first exchanges context among compact structural units. The de-tokenizer has 12 unit layers at width 768, followed by four atom-refinement layers at width 256. Each unit is expanded into its expected atoms:

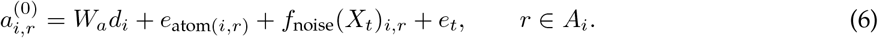

Here *d*_*i*_ combines the FSQ code and chemical features. A geometric atom refiner predicts a velocity for every atom. To resolve packing and interfaces, refinement includes bonded neighbors and dynamic inter-unit spatial neighbors, in addition to within-unit messages. Relative-coordinate geometric features and equivariant vector outputs preserve proper-rotation covariance.

### 3.2 Noise-to-structure training objective

We adopt a conditional flow-matching implementation of the generative de-tokenizer. After centering the assembly and scaling coordinates by a fixed 10 Å length unit, draw Gaussian noise *X*_0_ and *t* ~ U (0, 1), and define

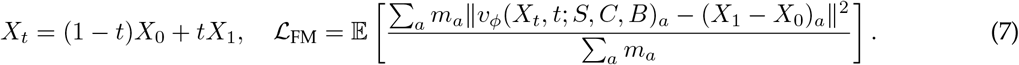

Because invariant tokens cannot specify an absolute frame, training uses rigid augmentation and an optimal proper rotation of the reference to the initial noise before interpolation. This alignment is a training coupling, not an inference input. An assembly is aligned jointly, never chain by chain; atom permutations are limited to chemical symmetries. At inference the ODE starts from centered noise and produces an arbitrary global pose. Predicted coordinates are rescaled to Å.

We supplement flow matching with chemical constraints evaluated on the endpoint estimate 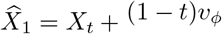:

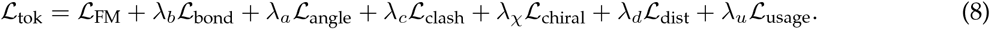

Bond and angle terms enforce chemistry; clash terms penalize nonbonded overlap; signed-volume terms preserve stereocenters; sampled intra- and intermolecular distance terms encourage fold and interface consistency. Losses are normalized by eligible atom or pair counts and fixed physical scales. Geometry terms are down-weighted near pure noise to avoid unstable endpoint extrapolation.

The utilization penalty acts on soft assignments to discrete levels, promoting batch diversity without forcing rare chemistries into an identical distribution.

### 3.3 Sampling

The inference budget is 50 Heun integration steps per sample (100 velocity evaluations). All chains and ligands are decoded jointly. Reconstruction uses reference-derived tokens, whereas prediction uses language-model-derived tokens.

### 3.4 From chemical input to a joint molecular structure

The language model receives all molecular identities and chemical connectivity before generating structural IDs. The de-tokenizer combines these IDs with atom identities and bonds to reconstruct the assembly jointly. Figure 3 shows chemical-unit granularity and confidence-based candidate selection; native coordinates are absent from the prediction inputs.

**Figure 3.**
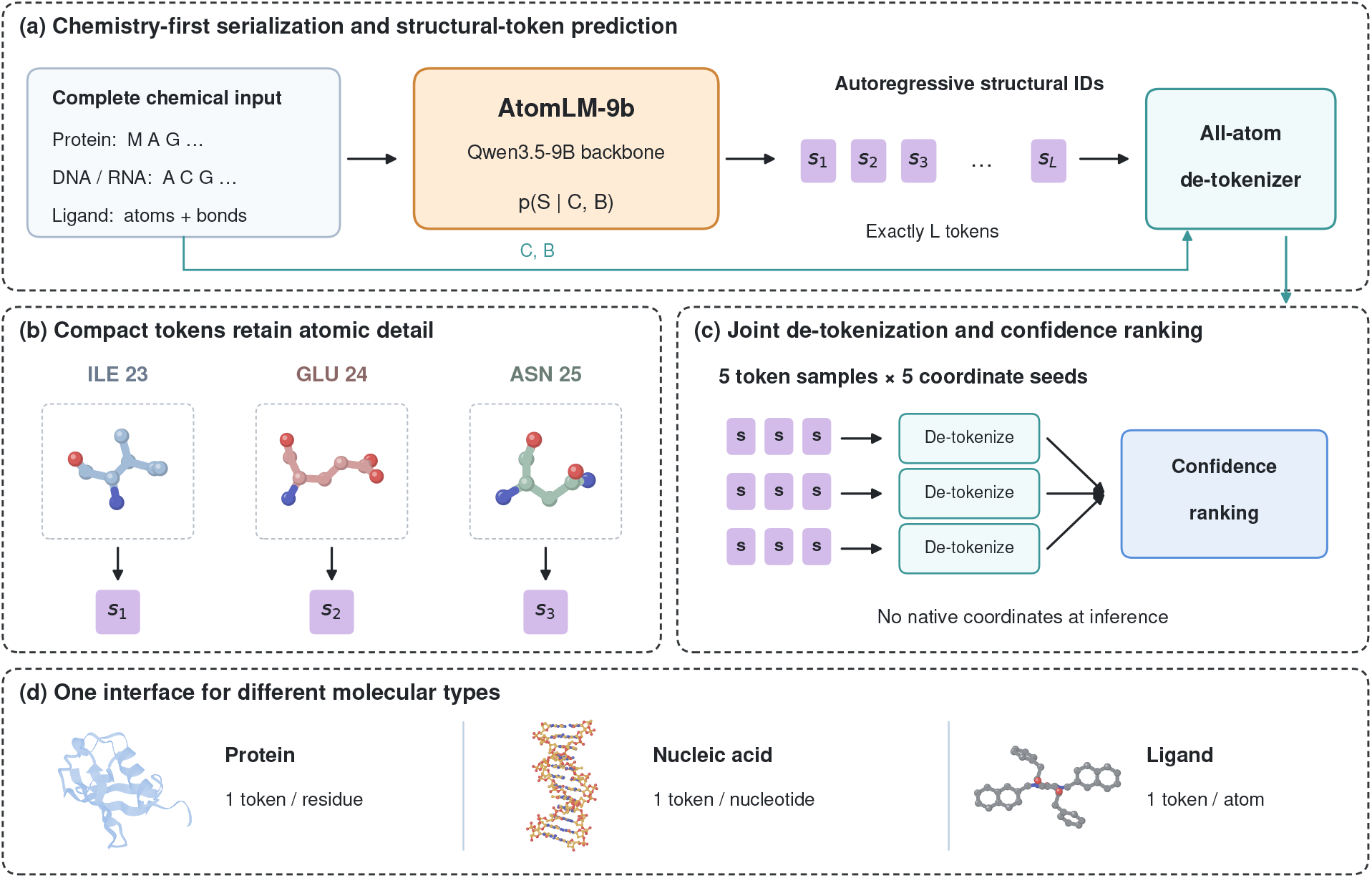
AtomLM-9b for structure prediction. (a) Given chemistry conditions a Qwen3.5-9B-based model; predicted structural IDs and chemical records condition joint decoding. (b) Ile23, Glu24, and Asn25 from 1UBQ chain A illustrate one token per protein residue. (c) The inference budget is five token samples and five coordinate seeds per sample; the three displayed rows only illustrate the workflow. Confidence ranking uses no native coordinates. (d) Protein, DNA, and ligand coordinate exemplars from 1UBQ, 1BNA, and XK2 in 1HVR illustrate token granularity.

## 4 Adapting Qwen3.5-9B to structural tokens

### 4.1 Backbone and vocabulary

AtomLM-9b starts from the language backbone of the released Qwen3.5-9B checkpoint. Its official text configuration specifies 32 hybrid layers with a 3:1 ratio of linear-attention to full-attention layers, hidden width 4,096, an FFN width of 12,288, and untied input and output embeddings. Full-attention layers use 16 query heads and four key/value heads [4]. We retain the causal language-model architecture and extend its 248,320-entry vocabulary with 8,192 structural-token IDs, dedicated chemical-unit symbols, and delimiters. Structural IDs are inserted directly into the vocabulary, rather than represented as decimal strings or split by text BPE.

With untied embeddings, structural symbols add 2 × 8192 × 4096 = 67,108,864 parameters; chemical and control symbols add a smaller amount. AtomLM-9b uses a backbone in the 9-billion-parameter class. The training context is 8,192 tokens, below the checkpoint’s configured 262,144-position limit.

### 4.2 Chemistry-first serialization

An input prefix lists the complete assembly before structure tokens are generated. Proteins and nucleic acids use one dedicated identity token per residue or nucleotide, with chain boundaries and stoichiometry. Ligands use canonical atom records followed by bond records containing indexed atom pairs, bond order, and stereochemistry. The atom ordering used for chemistry is also used for structural IDs. Covalent inter-chain and protein–ligand bonds are explicit. Equivalent chain permutations are randomized during training and fixed by a documented inference convention.

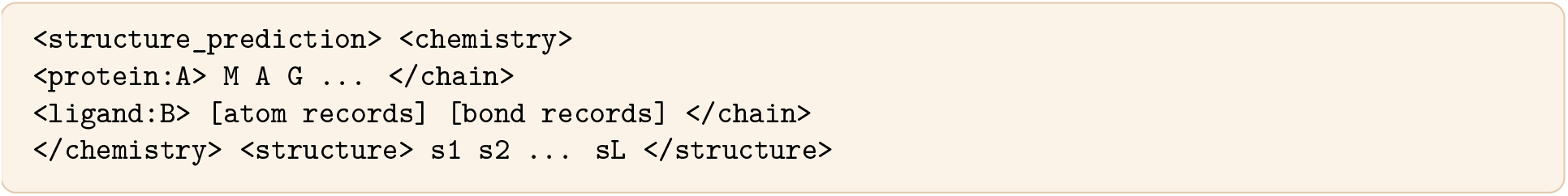

Chemical records determine output length and atom inventory. The model generates exactly one structural ID per unit, using a grammar mask over the structural vocabulary and a deterministic closing delimiter after *L* IDs. This guarantees format compliance, not accuracy. The primary mode uses a single sequence per chain, no MSA, no structural templates, and no native pocket coordinates.

### 4.3 Autoregressive structural learning

For fixed chemistry, the sequence model and coordinate de-tokenizer factorize prediction as

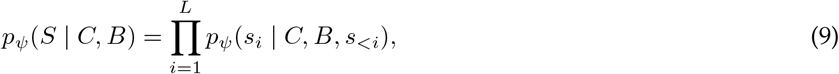

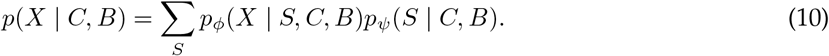

For chemistry-to-structure prediction, training uses teacher-forced structural cross-entropy, with loss masked on the chemical prefix. Language-model training additionally incorporates structure–text annotation pairs to align discrete structural representations with natural-language descriptions. The frozen tokenizer supplies labels; its version and code-to-ID map remain fixed throughout language-model training. Otherwise the same ID can change meaning across epochs.

### 4.4 Joint generation and confidence ranking

The inference budget is five token samples, each decoded with five coordinate-noise seeds: 25 candidates per target. Token sampling uses temperature 0.8 and top-*p* = 0.95, selected on validation data. A geometry-aware confidence network scores the assembly from chemistry, predicted coordinates, and model states. It is trained on out-of-fold predictions to estimate local accuracy, interface quality, and clashes. Native coordinates are never available to this ranker at inference. Top-ranked predictions are the primary endpoint. Language-model likelihood alone is not treated as calibrated structural confidence.

## 5 A staged training recipe

We train the system in five stages. The core dependency is sequential: establish a stable geometric vocabulary, freeze it, then train the structural language model. Complex supervision is necessary because monomer reconstruction alone does not teach relative binding poses.

### 5.1 Data sources and preprocessing

The tokenizer corpus combines experimental PDB biological assemblies, high-confidence predicted monomers, and small-molecule conformers with explicit bond graphs. The language-model corpus includes both chemistry–structure pairs and aligned structure–text annotation pairs. The latter associate biomolecular structures, represented by all-atom structural tokens, with their corresponding natural-language annotations, providing supervision for alignment between structural and textual representations. The collection is deduplicated, with experimental complexes receiving extra sampling weight. AFDB structures are teacher predictions and are tracked separately from experimental coordinates.

Preprocessing standardizes chain and atom identifiers, chooses a consistent alternate-location conformer, preserves biological assembly transforms, resolves ligand stereochemistry, and flags incomplete residues. Missing coordinates are loss-masked rather than set to zero. Random rigid rotations and chain-order per-mutations regularize serialization. Crops retain interfaces and full ligand graphs; later stages include complete assemblies so that crop-level accuracy does not conceal assembly failures.

The modality mixture is 50% protein monomers, 30% protein-containing complexes, 10% nucleic-acid-containing examples, and 10% small molecules, with mutually exclusive buckets (nucleic-acid-containing complexes enter the third bucket). These percentages describe molecular composition; text annotations are paired with the corresponding structures rather than defining a separate molecular category. Within complex buckets, antibody, ligand, and ion interfaces are balanced. Per-modality validation curves guide training. Length bucketing limits padding, while target-level normalization prevents large complexes from dominating every update.

### 5.2 Optimization and systems considerations

We train AtomLM-9b on 64 NVIDIA H100 GPUs for one week. We use BF16 training, AdamW (*β*_1_, *β*_2_) = (0.9, 0.95), weight decay 0.1, gradient clipping at 1.0, 3% warmup, and cosine decay. Each language-model batch contains approximately 2^20^ non-padding tokens per optimizer update through distributed accumulation. Gradient checkpointing and sharded optimizer states control memory. Token caches record tokenizer hashes, chemical ordering, and coordinate provenance. Packed sequences use attention isolation between assemblies.

### 5.3 Checkpoint selection

Tokenizer checkpoints are selected using reconstruction loss. Language-model checkpoints use validation token loss together with decoded accuracy; better perplexity need not imply better geometry. The ranker uses disjoint validation candidates. Benchmark targets are never used to choose learning rates, sampling temperature, de-tokenizer steps, or checkpoints.

## 6 Evaluation protocol

We evaluate tokenizer reconstruction and end-to-end structure prediction separately. Reconstruction uses reference-derived structural tokens and reports protein C*α*-lDDT, all-atom lDDT, all-atom RMSD, and TM-score, together with small-molecule conformer RMSD. Structure prediction generates tokens from chemical inputs without reference coordinates.

### FoldBench reference

AlphaFold 3 (AF3) [5] values are taken directly from Supplementary Table 3 of the FoldBench paper [6], using the full benchmark rather than the post-January-2024 subset. The official leaderboard provides the corresponding interface and nucleic-acid results [7]. Monomers are scored by mean lDDT. Protein–ligand success requires ligand RMSD *<* 2 Å and lDDT-PLI *>* 0.8; other interface tasks use DockQ ≥ 0.23. We report results on eight FoldBench tasks.

**Table 1.** Token granularity. One structural ID is emitted per unit; chemical identities and connectivity are separate inputs.

| Entity | Structural unit | Geometry recovered |
|---|---|---|
| Protein / peptide | One amino-acid residue | Backbone and side-chain heavy atoms |
| DNA / RNA | One nucleotide | Base, sugar, and phosphate atoms |
| Small molecule | One heavy atom | Joint conformation and bound pose |
| Ion | One ion | Position and interaction context |
| Modified residue | CCD-defined residue or atom fallback | Explicit chemical graph |

**Table 2.**
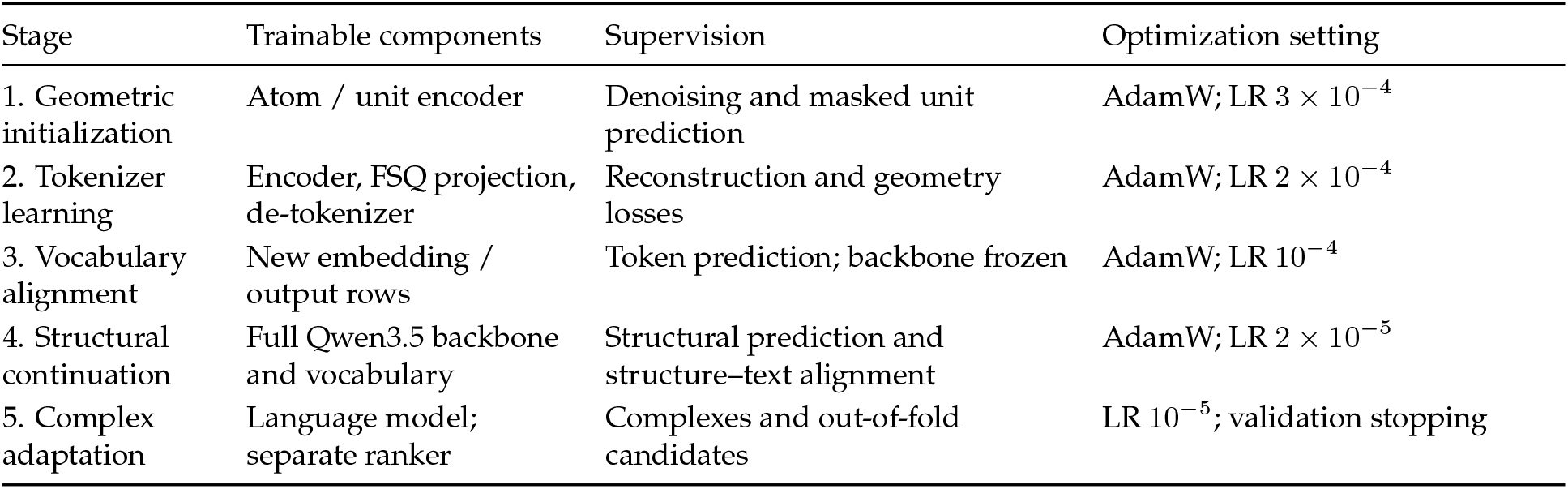
Training stages and optimization settings.

**Table 3.** All-Atom Structure Tokenizer reconstruction on proteins. lDDT and TM-score are on [0, 1]; RMSD is in Å.

| Dataset | Cα-lDDT↑ | AA-lDDT↑ | AA-RMSD↓ | TM-score↑ |
|---|---|---|---|---|
| CASP14 | 0.9020 | 0.8320 | 1.2500 | 0.9720 |
| CASP15 | 0.9010 | 0.8350 | 1.7400 | 0.9510 |

### Prediction settings

AtomLM-9b uses single-sequence, template-free inputs and selects the highest-ranked structure among 25 candidates (five token samples, each with five coordinate seeds). AF3 entries are published reference results obtained with the FoldBench inference pipeline and its sequence resources; they are not AF3 reruns in this study.

### Data separation

Structural training uses a PDB release cutoff of January 12, 2023, with benchmark-specific homology and chemical-similarity filtering. CASP14/15 reconstruction excludes evaluation targets and their homologs from its training data. Qwen3.5’s general pretraining corpus is not fully auditable.

## 7 Tokenizer reconstruction

### 7.1 Protein reconstruction

At one structural token per residue, the tokenizer achieves all-atom lDDT of 0.832 on CASP14 and 0.835 on CASP15 (Table 3). Corresponding all-atom RMSDs are 1.25 and 1.74 Å, with TM-scores of 0.972 and 0.951.

### 7.2 Small-molecule reconstruction

Small-molecule reconstruction yields RMSDs of 0.0330–0.0495 Å across QM9 and the three ∇^2^DFT splits (Table 4). These self-aligned reconstruction metrics describe internal conformation, rather than receptor-relative docking accuracy.

**Table 4.** All-Atom Structure Tokenizer reconstruction on small molecules.

| Dataset / split | RMSD (Å)↓ |
|---|---|
| QM9 | 0.0330 |
| ∇<sup>2</sup>DFT / conformer | 0.0480 |
| ∇<sup>2</sup>DFT / structure | 0.0490 |
| ∇<sup>2</sup>DFT / scaffolds | 0.0495 |

## 8 Structure-prediction benchmark

Table 5 compares AtomLM-9b with the published AF3 full-set results on FoldBench. AtomLM-9b achieves protein-monomer lDDT of 0.858 versus 0.88 for AF3, a difference of −0.022. The remaining results show task-specific differences rather than uniform agreement across molecular types.

**Table 5.** FoldBench structure prediction. AF3 values are reproduced from Supplementary Table 3 of the FoldBench paper [6]. Δ is AtomLM-9b minus AF3, calculated from the displayed values: absolute lDDT for monomers and percentage points for interfaces.

| Task | Metric↑ | AF3 (published) | AtomLM-9b | Δ |
|---|---|---|---|---|
| Protein monomer | lDDT | 0.88 | 0.858 | −0.022 |
| RNA monomer | lDDT | 0.61 | 0.700 | +0.090 |
| DNA monomer | lDDT | 0.53 | 0.774 | +0.244 |
| Protein–protein | Success (%) | 72.93 | 75.01 | +2.08 |
| Antibody–antigen | Success (%) | 47.90 | 44.00 | −3.90 |
| Protein–ligand | Success (%) | 64.90 | 67.45 | +2.55 |
| Protein–RNA | Success (%) | 62.32 | 63.21 | +0.89 |
| Protein–DNA | Success (%) | 79.18 | 78.86 | −0.32 |

AtomLM-9b is higher on RNA and DNA monomer lDDT and on protein–protein, protein–ligand, and protein–RNA success, while AF3 is higher on protein monomers, antibody–antigen, and protein–DNA pre-diction. These comparisons describe the reported point estimates.

## 9 Ablations and limitations

### 9.1 Ablation results

Table 6 reports diagnostic comparisons using the CASP15 and protein–ligand endpoints. Each architectural variant was trained under the same protocol, with matched data and optimization budgets. The comparisons isolate encoder, quantizer, de-tokenizer, and complex-adaptation contributions.

**Table 6.**
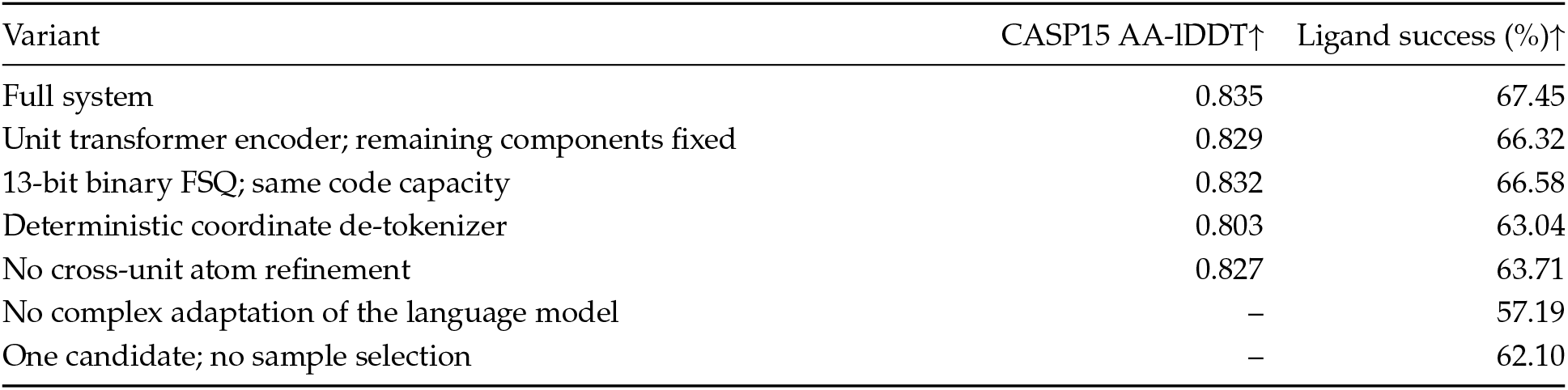
Ablation results. A dash denotes an endpoint not varied by that ablation.

| Variant | CASP15 AA-lDDT↑ | Ligand success (%)↑ |
|---|---|---|
| Full system | 0.835 | 67.45 |
| Unit transformer encoder; remaining components fixed | 0.829 | 66.32 |
| 13-bit binary FSQ; same code capacity | 0.832 | 66.58 |
| Deterministic coordinate de-tokenizer | 0.803 | 63.04 |
| No cross-unit atom refinement | 0.827 | 63.71 |
| No complex adaptation of the language model | – | 57.19 |
| One candidate; no sample selection | – | 62.10 |

Replacing the hierarchical encoder reduces CASP15 AA-lDDT from 0.835 to 0.829 and ligand success from 67.45% to 66.32%. Removing cross-unit atom refinement reduces ligand success to 63.71%, while omitting complex adaptation reduces it to 57.19%. Using one candidate yields 62.10% success; this row measures the effect of candidate selection rather than an architectural change.

### 9.2 Technical limitations

A finite structural vocabulary compresses side-chain and interface information, and autoregressive token errors propagate through an assembly. Rare chemical components and complex binding geometries remain challenging. The model represents heavy atoms; hydrogens and protonation-dependent interactions are outside the present scope. Reconstruction and free-running structure prediction remain distinct tasks.

## 10 Conclusion

Scinetics AtomLM-9b combines an interaction-aware encoder, FSQ structural tokens, a joint generative coordinate de-tokenizer, and a Qwen3.5-9B structural language model. The all-atom tokenizer achieves high-fidelity reconstruction through a compact discrete representation of biomolecular geometry. Built on this representation, the trained structural language model achieves protein-monomer lDDT of 0.858 on FoldBench, approaching the published AF3 reference of 0.88. Together, these results demonstrate that discrete all-atom representations support both accurate atomic reconstruction and competitive language-model-based structure prediction.

## References

[1] A. Fang, M. Desgagné, Z. Zhang, A. Zhou, J. Loscalzo, B. L. Pentelute, and M. Zitnik. Learning Universal Representations of Intermolecular Interactions with ATOMICA. bioRxiv, supplied v4, September 7, 2026. doi:10.1101/2025.04.02.646906. Architecture: Methods 1.3 and 2.

[2] F. Mentzer, D. Minnen, E. Agustsson, and M. Tschannen. Finite Scalar Quantization: VQ-VAE Made Simple. ICLR, 2024. https://arxiv.org/abs/2309.15505.

[3] Qwen Team. Qwen3.5-9B: official model card. Accessed October 27, 2026. https://huggingface.co/Qwen/Qwen3.5-9B.

[4] Qwen Team. Qwen3.5-9B: official model configuration. Accessed October 27, 2026. https://huggingface.co/Qwen/Qwen3.5-9B/blob/main/config.json.

[5] J. Abramson et al. Accurate structure prediction of biomolecular interactions with AlphaFold 3. Nature 630, 493–500, 2024. doi:10.1038/s41586-024-07487-w.

[6] S. Xu, Q. Feng, L. Qiao, H. Wu, T. Shen, Y. Cheng, S. Zheng, and S. Sun. Benchmarking all-atom biomolecular structure prediction with FoldBench. Nature Communications 17, 442 (2026); online December 4, 2025. doi:10.1038/s41467-025-67127-3.

[7] BEAM Labs. FoldBench: official repository and evaluation protocol. Accessed October 27, 2026. https://github.com/BEAM-Labs/FoldBench.

[8] X. Huang, E. Su, A. Wang, P. Tan, S. Xu, Y. Wang, C. Tang, S. Tang, and M. Li. ADT: All-Atom Diffusion Tokenizer for 3D Molecular Structures. arXiv:2610.23265, 2026. https://arxiv.org/abs/2610.23265.

